# Pancreatic cancer disrupts the adult hippocampal neurogenic niche

**DOI:** 10.64898/2026.07.03.736329

**Authors:** Dimitrios Troumpoukis, Adriana Papadimitropoulou, Chrysanthi Charalampous, Paraskevi Kogionou, Alexia Polissidis, Nicolas Nicolaides, Yassemi Koutmani, Ioannis Serafimidis

## Abstract

Pancreatic cancer (PC) exhibits a striking association with depression, with neuropsychiatric symptoms frequently preceding diagnosis. However, the biological mechanisms linking pancreatic tumor development to central nervous system dysfunction remain poorly understood. Here, we investigated the impact of PC progression on adult hippocampal neurogenesis using complementary orthotopic xenograft and genetically engineered mouse models. Tumor-bearing mice developed depressive-like behavioral abnormalities accompanied by reduced adult hippocampal neurogenesis, including depletion of neural stem cell populations and immature neurons in both dorsal and ventral dentate gyrus regions. In the genetic model, neurogenic impairment progressed in parallel with disease severity. Exposure of primary hippocampal neural stem cells to serum derived from tumor-bearing mice selectively impaired cell survival, indicating that circulating factors are sufficient to compromise neurogenic capacity. Consistent with this, cytokine profiling revealed profound systemic inflammatory alterations, with IL-6 emerging as the only cytokine consistently elevated across both models. Together, our findings identify disruption of the adult hippocampal neurogenic niche as a previously unrecognized consequence of pancreatic cancer progression and provide a biological framework for pancreatic cancer-associated depression.

## INTRODUCTION

Pancreatic cancer (PC) is among the most aggressive human malignancies, characterized by rapid progression, early metastatic dissemination, lack of effective early detection, and poor response to currently available therapies ^1^. As a result, prognosis remains dismal. Beyond its devastating oncological course, PC is also notable for its strong association with neuropsychiatric symptoms, particularly major depression. Among tumors of the digestive system, PC shows the highest incidence of depression, and depressive symptoms often emerge months before formal cancer diagnosis, a phenomenon commonly referred to as Pancreatic Cancer-Associated Depression (PCAD) ^2^. Depressive symptoms have been reported in up to 50% of patients with PC and are frequently accompanied by anxiety, insomnia, and fatigue ^3–11^. Importantly, depression in PC is often refractory to treatment ^12^ and has been associated with worse clinical outcome, including reduced 5-year survival ^13,14^. Despite this striking clinical association, the biological mechanisms linking pancreatic tumor development to depression remain poorly understood.

One of the processes that underlie depressive-like phenotypes and is highly modulated by systemic alterations, providing a possible mechanistic link between cancer progression and affective dysfunction, is adult hippocampal neurogenesis. In the adult mammalian brain, new neurons continue to be generated from neural stem cells (NSCs) in the subgranular zone (SGZ) of the dentate gyrus (DG), a specialized neurogenic niche within the hippocampus ^15–18^. Adult hippocampal neurogenesis contributes to cognitive and emotional functions, including pattern separation, contextual discrimination, memory, and mood regulation ^19^. A large body of work has linked impaired hippocampal neurogenesis to depressive states, and several animal models of depression exhibit reduced neurogenesis across the hippocampal axis ^20–22^. In some cases, these alterations appear particularly prominent in the ventral hippocampus, a region more closely associated with emotional behavior, whereas the dorsal hippocampus is classically linked to learning and memory ^23^. Together, these observations raise the possibility that disrupted adult hippocampal neurogenesis may contribute to the neuropsychiatric manifestations of PC.

Adult neurogenesis is constantly regulated by factors from the local and systemic environment. Within the hippocampal niche, NSCs are sensing signals derived by the neighboring astrocytes, microglia, vasculature, and neurons, and respond adequately by altering their behavior, stem cell quiescence, proliferation, differentiation, and survival ^24–26^. At the same time, NSCs remain highly responsive to peripheral cues. Circulating inflammatory cytokines, stress-related mediators, and metabolic factors can influence the neurogenic niche either by crossing the blood-brain barrier under defined conditions or by signaling indirectly through endothelial, immune, or neural pathways. Under pathological conditions, these systemic signals often suppress neurogenesis by reducing NSC survival and proliferative activity ^27^. This is especially relevant in cancer, where tumor progression is frequently accompanied by profound systemic inflammation.

Thus, in In pancreatic cancer, inflammatory mediators have long been implicated in disease progression and symptom burden. Notably, a small but important body of clinical evidence suggests that cytokine dysregulation may contribute specifically to PCAD. In particular, patients with PC who develop depression display elevated circulating IL-6 levels compared to non-depressed patients with PC and to healthy controls ^28,29^. These findings raise the possibility that systemic inflammatory signals, and IL-6 in particular, may participate in the communication between pancreatic tumors and the brain. However, whether such factors affect hippocampal neurogenesis during PC progression, and whether they contribute to depressive-like phenotypes, remains unknown.

Here, we investigated the impact of pancreatic tumor development on adult hippocampal neurogenesis using complementary xenograft and genetic mouse models of PC. We combined behavioral analysis with histological and immunofluorescence-based assessment of the hippocampal neurogenic lineage, examined the direct effects of tumor-derived serum on hippocampal NSC proliferation and survival in vitro, and profiled systemic inflammatory mediators to identify candidate factors associated with the observed phenotype. Our findings identify impaired adult hippocampal neurogenesis as a feature of pancreatic cancer progression and support a model in which systemic inflammatory signals, with IL-6 emerging as a shared candidate mediator across models, contribute to the tumor–brain crosstalk underlying pancreatic cancer-associated depressive-like phenotypes.

## RESULTS

### NOD-SCID mice bearing pancreatic tumors display depressive-like behavior

To investigate the impact of pancreatic cancer (PC) on central nervous system (CNS) function, we established an orthotopic xenograft mouse model by injecting human Panc-1 cells into the pancreas of immunocompromised NOD-SCID mice (hereafter referred to as Xeno^Panc1^) (Fig. 1A). This approach resulted in reproducible tumor formation with defined growth kinetics. Histological analysis of pancreatic sections at 5-, 11- and 21-days post-injection (p.i.) confirmed progressive tumor development (Fig. 1B). More specifically, at 5 days p.i., tumor and normal pancreatic tissue were not readily distinguishable. By 11 days p.i., well-defined tumor masses were evident, whereas at 21 days p.i., tumors had extensively infiltrated the pancreas, were macroscopically detectable, and in some cases showed signs of metastasis. Based on these observations, we defined 9–11 days p.i. as an early stage of disease and performed behavioral analyses within this time window (Fig. 1C). To assess locomotor activity mice were subjected to the open field test (OFT) while anxiety and depression-related behavior was evaluatedby elevated plus maze (EPM), and tail suspension test (TST). In the OFT, no significant differences were observed between Xeno^Panc1^ and sham-operated, Xeno^Sham^ (c), mice in either total distance traveled or time spent in the center of the arena (Fig. S1A), indicating preserved general locomotor activity and exploratory behavior. Similarly, total distance traveled in the EPM was comparable between groups (Fig. 1D), suggesting that tumor-bearing mice did not exhibit locomotor impairments or general sickness behavior that could confound behavioral outcomes. However, Xeno^Panc1^ mice spent significantly less time in the open arms of the EPM (Fig. 1D). In addition, in the TST, tumor-bearing mice exhibited a marked increase in immobility time compared to controls (Fig. 1E). Together, these findings indicate the emergence of a depressive-like behavioral phenotype in mice during the early stages of pancreatic tumor development.

**Fig 1.**
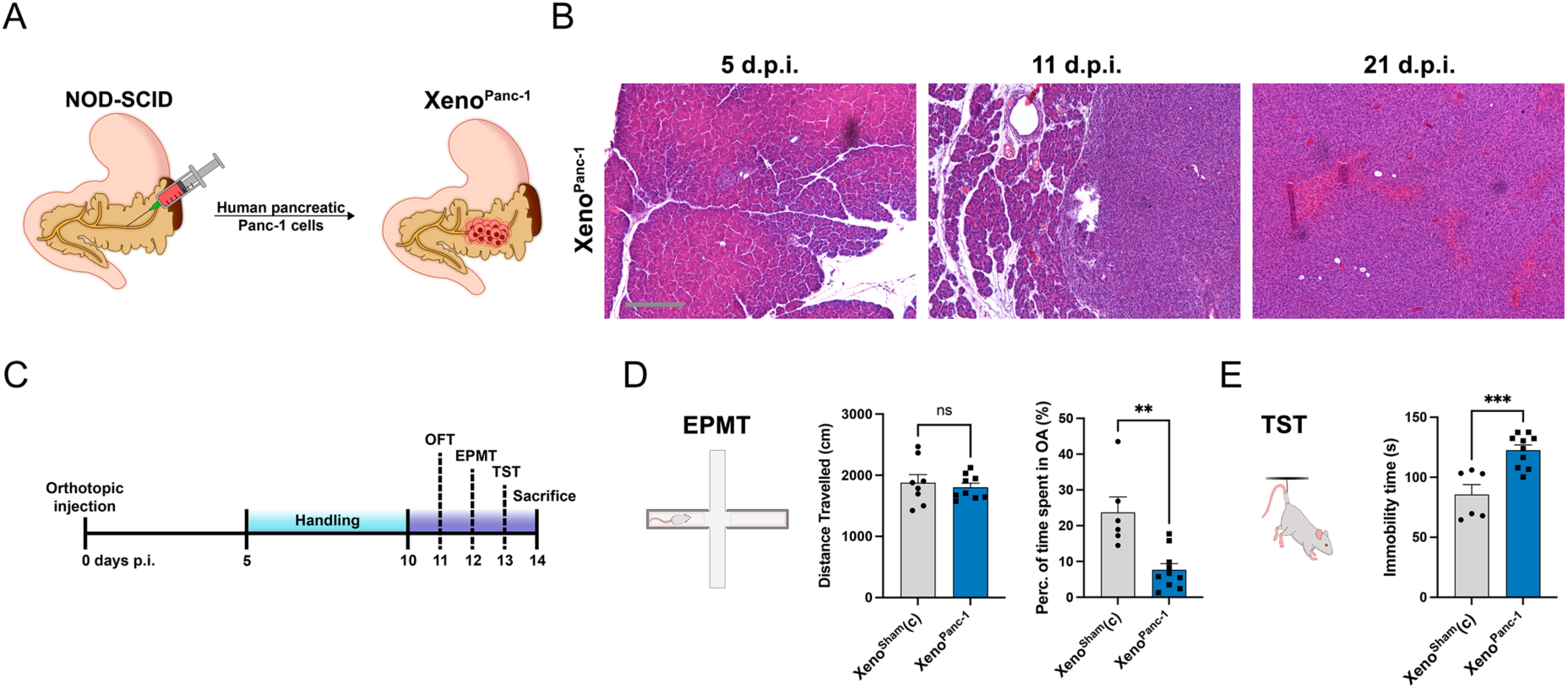
Xeno^Panc-1^ mice exhibit depressive-like behavior. A) Schematic representation of the generation of Xeno^Panc-1^. B) Representative images of pancreatic tissue stained with H&E at 5-, 11- and 21-days post-injection with human pancreatic cancer cells. Scale bar 200μm. C) Schematic diagram of the behavioral experimental timeline. D) Quantification of mobility (distance travelled in the arena) and anxiety-phenotype (time spent in Open Arms) in the Elevated Plus Maze Test. n=8 Xeno^Sham(c)^ mice and n=10 Xeno^Panc-1^ mice. Data were analyzed with the two-tailed unpaired student’s t-test. Error bars represent SEM. E) Quantification of Immobility time in the Tail Suspension Test. n=8 Xeno^Sham(c)^ mice and n=10 Xeno^Panc-1^ mice. Data were analyzed with the Mann-Whitney test. Error bars represent SEM.

### Adult hippocampal neurogenesis is impaired in NOD-SCID mice bearing pancreatic tumors

Since adult neurogenesis in the dentate gyrus (DG) of the hippocampus is linked to depressive-like behaviour, we sought to investigate the neurogenic activity in the hippocampus of Xeno^Panc1^ mice and the involvement of this process in the observed depressive-like phenotype. More specifically, brains were collected from tumor-bearing and control mice one day after completion of behavioral testing and proceeded to immunofluorescence analysis using specific antibodies against established markers of the neurogenic lineage (Fig. S2A). Thus, to detect neural stem cells (NSCs) in an either quiescent or early-activated state, expression of the marker GFAP in combination with morphological criteria was used, while proliferative activity and commitment of NSCs to the neuronal lineage were assessed using antibodies against SOX2 and doublecortin (DCX) respectively (Fig. S2A).

Given the functional heterogeneity of the hippocampus, we analyzed separately the dorsal and ventral regions of the DG. The dorsal DG is primarily associated with cognitive functions such as learning and memory, whereas the ventral DG is more closely linked to mood regulation and emotional behavior. Quantitative analysis revealed a marked reduction in neurogenesis in Xeno^Panc1^ mice. In the dorsal DG, the number of DCX⁺ immature neurons was significantly decreased compared to sham-operated controls (Fig. 2A, B). In parallel, the NSC pool was also reduced, as indicated by a lower number of GFAP⁺ radial glia-like (RGL) cells, which represent the self-renewing NSC population in the adult hippocampus (Fig. 2C, D). Consistently, SOX2⁺ cells were also significantly decreased (Fig. E, F), along with a reduction in SOX2⁺/DCX⁺ double-positive cells (Fig. G, H), suggesting impaired progression along the neurogenic lineage.

**Fig 2.**
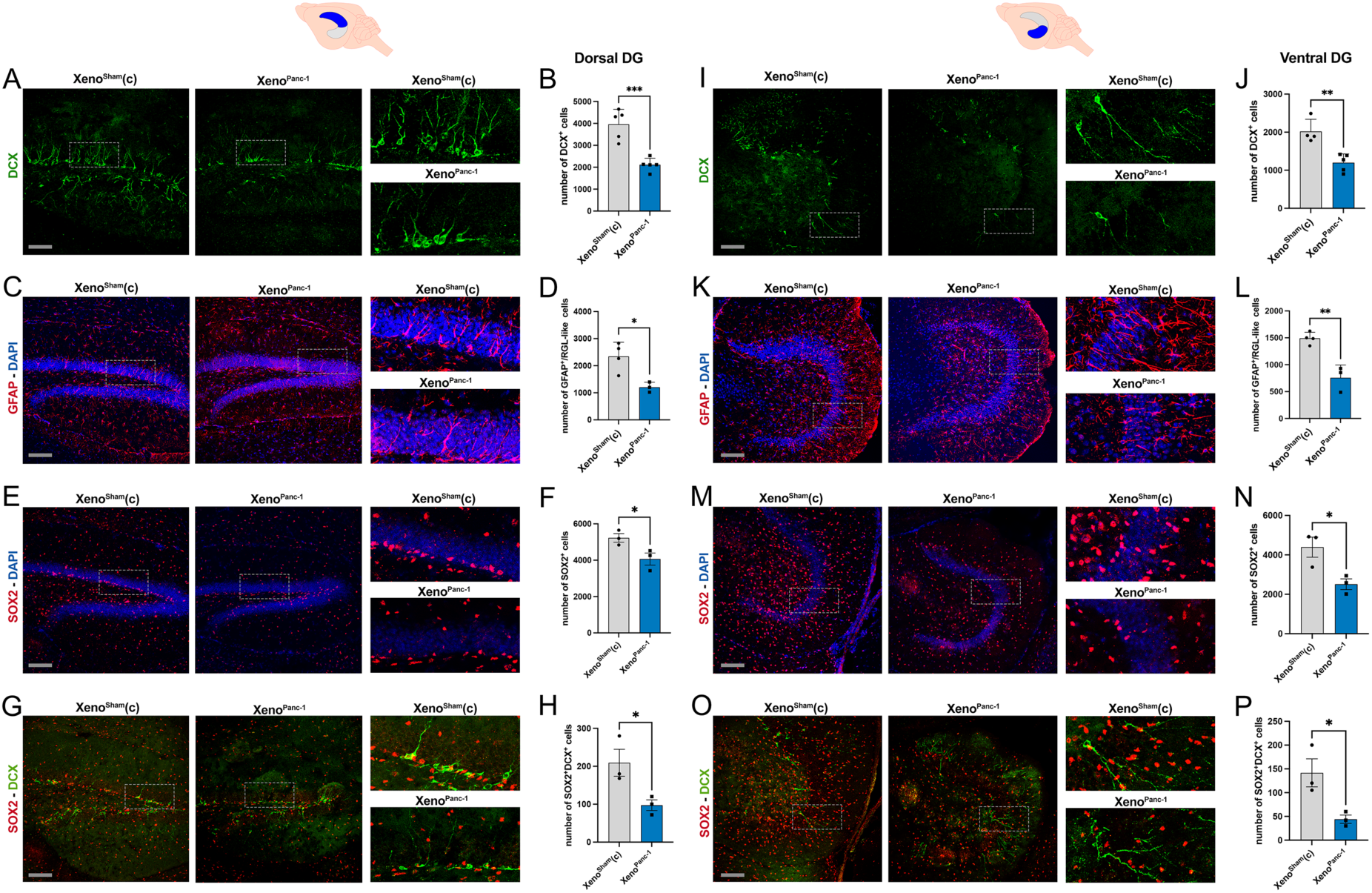
Adult hippocampal neurogenesis is impaired in Xeno^Panc-1^ mice. A) Representative images of DCX^+^ cells in the dorsal DG of Xeno^Sham(c)^ and Xeno^Panc-1^ mice. Scale bar = 200μm. B) Quantification of numbers of DCX⁺ cells in the dorsal DG of Xeno^Sham(c)^ and Xeno^Panc-1^ mice. n=5 mice for each group. Data were analyzed with the two-tailed unpaired student’s t-test. Error bars represent SEM. C) Representative images of GFAP^+^ cells in the dorsal DG of Xeno^Sham(c)^ and Xeno^Panc-1^ mice. DAPI was used as counterstain. Scale bar = 200μm. D) Quantification of numbers of GFAP⁺ cells in the dorsal DG of Xeno^Sham(c)^ and Xeno^Panc-1^ mice. n=4 Xeno^Sham(c)^ mice and n=3 Xeno^Panc-1^ mice. Data were analyzed with the two-tailed unpaired student’s t-test. Error bars represent SEM. E) Representative images of SOX2^+^ cells in the dorsal DG of Xeno^Sham(c)^ and Xeno^Panc-1^ mice. DAPI was used as counterstain. Scale bar = 200μm. F) Quantification of numbers of SOX2⁺ cells in the dorsal DG of Xeno^Sham(c)^ and Xeno^Panc-1^ mice. n=3 for each group. Data were analyzed with the two-tailed unpaired student’s t-test. Error bars represent SEM. G) Representative images of SOX2^+^DCX^+^ cells in the dorsal DG of Xeno^Sham(c)^ and Xeno^Panc-1^ mice. Scale bar = 200μm. H) Quantification of numbers of SOX2^+^DCX⁺ cells in the dorsal DG of Xeno^Sham(c)^ and Xeno^Panc-1^ mice. n=3 for each group. Data were analyzed with the two-tailed unpaired student’s t-test. Error bars represent SEM. I) Representative images of DCX^+^ cells in the ventral DG of Xeno^Sham(c)^ and Xeno^Panc-1^ mice. Scale bar = 200μm. J) Quantification of numbers of DCX⁺ cells in the ventral DG of Xeno^Sham(c)^ and Xeno^Panc-1^ mice. n=4 Xeno^Sham(c)^ mice and n=5 Xeno^Panc-1^ mice. Data were analyzed with the two-tailed unpaired student’s t-test. Error bars represent SEM. K) Representative images of GFAP^+^ cells in the ventral DG of Xeno^Sham(c)^ and Xeno^Panc-1^ mice. DAPI was used as counterstain. Scale bar = 200μm. L) Quantification of numbers of GFAP⁺ cells in the ventral DG of Xeno^Sham(c)^ and Xeno^Panc-1^ mice. n=4 Xeno^Sham(c)^ mice and n=3 Xeno^Panc-1^ mice. Data were analyzed with the two-tailed unpaired student’s t-test. Error bars represent SEM. M) Representative images of SOX2^+^ cells in the ventral DG of Xeno^Sham(c)^ and Xeno^Panc-1^ mice. DAPI was used as counterstain. Scale bar = 200μm. N) Quantification of numbers of SOX2⁺ cells in the ventral DG of Xeno^Sham(c)^ and Xeno^Panc-1^ mice. n=3 for each group. Data were analyzed with the two-tailed unpaired student’s t-test. Error bars represent SEM. O) Representative images of SOX2^+^DCX^+^ cells in the ventral DG of Xeno^Sham(c)^ and Xeno^Panc-1^ mice. Scale bar = 200μm. P) Quantification of numbers of SOX2^+^DCX⁺ cells in the ventral DG of Xeno^Sham(c)^ and Xeno^Panc-1^ mice. n=3 for each group. Data were analyzed with the two-tailed unpaired student’s t-test. Error bars represent SEM.

Importantly, similar alterations were observed in the ventral DG. Xeno^Panc1^ mice exhibited a significant reduction in DCX⁺, GFAP⁺RGL, and SOX2⁺ cell populations in this region, indicating that neurogenesis is broadly impaired across hippocampal subregions (Fig. 2I-P). Although the ratio of SOX2⁺/DCX⁺ cells to total SOX2⁺ cells that depicts the survival and neuronal differentiation capacity of the SOX2+ neural stem cell population showed a decreasing trend in Xeno^Panc1^ mice in both dorsal and ventral DG, these differences did not reach statistical significance (Fig. S2B, C), suggesting that the primary effect may reflect a reduction in the overall NSC pool rather than a specific block in lineage progression.

Given the well-established role of insulin as a trophic factor supporting adult neurogenesis, we also assessed whether systemic insulin deficiency could account for the observed phenotype. ELISA measurements of circulating insulin levels revealed no significant differences between Xeno^Panc1^ and control mice (Fig. S2D), arguing against a major contribution of impaired insulin signaling.

Together, these results demonstrate that pancreatic tumor development is associated with a robust reduction in adult hippocampal neurogenesis, affecting both stem cell maintenance and neuronal differentiation, independent of systemic insulin levels.

### Impaired adult hippocampal neurogenesis is recapitulated in a genetic model of pancreatic cancer

To validate our findings in a physiologically relevant setting, we employed a genetically engineered mouse model of pancreatic ductal adenocarcinoma (PDAC), Pdx1^Cre^;Actin-Kras^G12D^ (hereafter referred to as Pdx1Cre-AKras*). In this model, pancreas-specific expression of oncogenic Kras^G12D^ under the constitutively active β-actin promoter, drives a stagewise and rapid progression to invasive PDAC, as previously described ^30^. Mice develop low-grade pancreatic intraepithelial neoplasias (PanINs) around postnatal day 10 (P10) and exhibit a median survival of approximately 45 days ^30^. Histological analysis confirmed progressive tumor development, with early lesions at P21 and more advanced disease at P40 (Fig. 3A,B).

**Fig 3.**
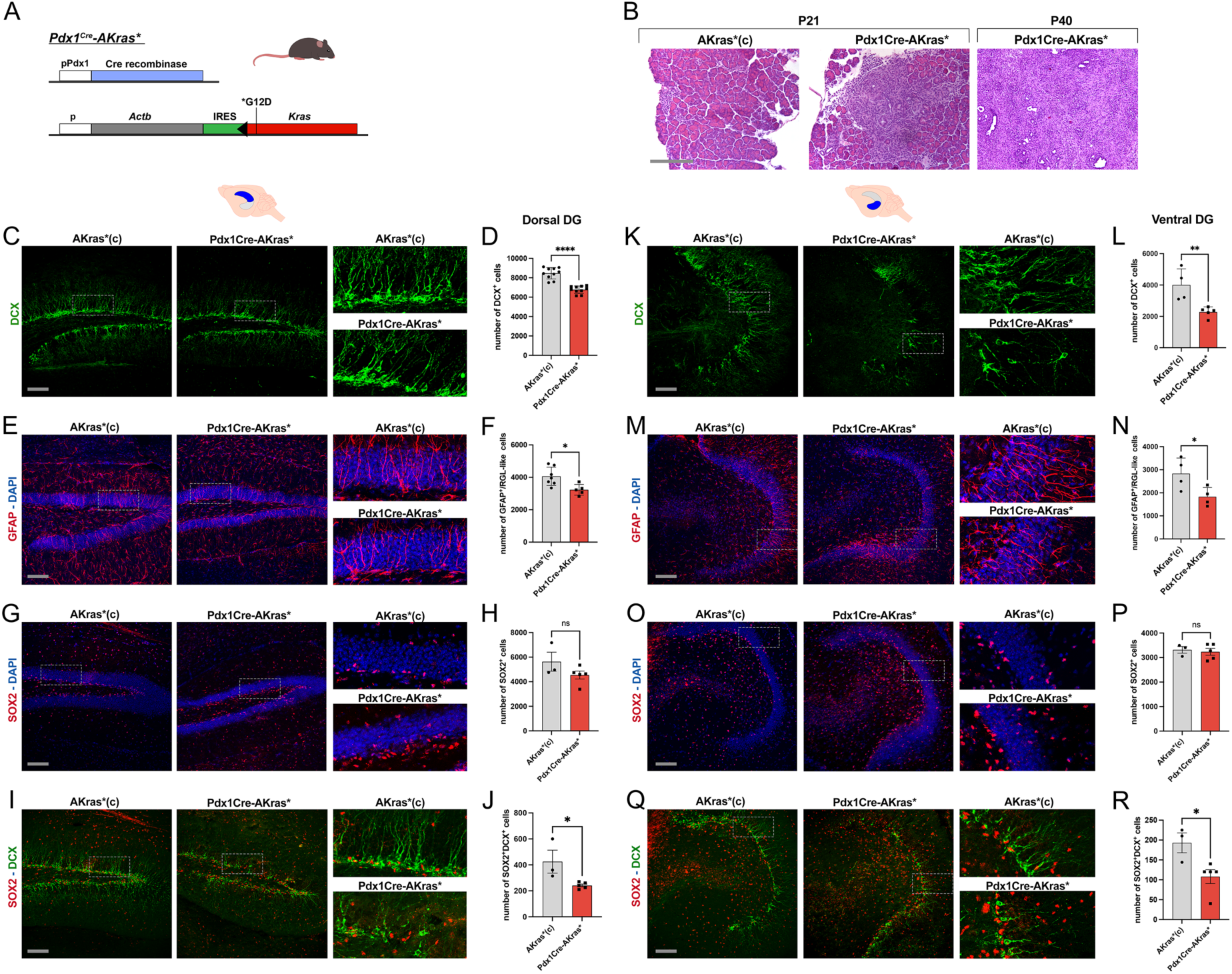
Adult Hippocampal Neurogenesis is impaired in Pdx1Cre-AKras* mice. A) Schematic representation of the Pdx1Cre-AKras* genotype. B) Representative images of pancreatic tissue stained with H&E at P21 and P40. Scale bar=200μm. C) Representative images of DCX^+^ cells in the dorsal DG of AKras*(c) and Pdx1Cre-AKras* mice. Scale bar = 200μm. D) Quantification of numbers of DCX⁺ cells in the dorsal DG of AKras*(c) and Pdx1Cre-AKras* mice. n=10 mice for each group. Data were analyzed with the two-tailed unpaired student’s t-test. Error bars represent SEM. E) Representative images of GFAP^+^ cells in the dorsal DG of AKras*(c) and Pdx1Cre-AKras* mice. DAPI was used as counterstain. Scale bar = 200μm. F) Quantification of numbers of GFAP⁺ cells in the dorsal DG of AKras*(c) and Pdx1Cre-AKras* mice. n=7 AKras*(c) mice and n=5 Pdx1Cre-AKras* mice. Data were analyzed with the two-tailed unpaired student’s t-test. Error bars represent SEM. G) Representative images of SOX2^+^ cells in the dorsal DG of AKras*(c) and Pdx1Cre-AKras* mice. DAPI was used as counterstain. Scale bar = 200μm. H) Quantification of numbers of SOX2⁺ cells in the dorsal DG of AKras*(c) and Pdx1Cre-AKras* mice. n=3 AKras*(c) mice and n=5 Pdx1Cre-AKras* mice. Data were analyzed with the two-tailed unpaired student’s t-test. Error bars represent SEM. I) Representative images of SOX2^+^DCX^+^ cells in the dorsal DG of AKras*(c) and Pdx1Cre-AKras* mice. Scale bar = 200μm. J) Quantification of numbers of SOX2^+^DCX⁺ cells in the dorsal DG of AKras*(c) and Pdx1Cre-AKras* mice. n=3 AKras*(c) mice and n=5 Pdx1Cre-AKras* mice. Data were analyzed with the two-tailed unpaired student’s t-test. Error bars represent SEM. K) Representative images of DCX^+^ cells in the ventral DG of AKras*(c) and Pdx1Cre-AKras* mice. Scale bar = 200μm. L) Quantification of numbers of DCX⁺ cells in the ventral DG of AKras*(c) and Pdx1Cre-AKras* mice. n=4 AKras*(c) mice and n=5 Pdx1Cre-AKras* mice. Data were analyzed with the two-tailed unpaired student’s t-test. Error bars represent SEM. M) Representative images of GFAP^+^ cells in the ventral DG of AKras*(c) and Pdx1Cre-AKras* mice. DAPI was used as counterstain. Scale bar = 200μm. N) Quantification of numbers of GFAP⁺ cells in the ventral DG of AKras*(c) and Pdx1Cre-AKras* mice. n=4 for each group. Data were analyzed with the two-tailed unpaired student’s t-test. Error bars represent SEM. O) Representative images of SOX2^+^ cells in the ventral DG of AKras*(c) and Pdx1Cre-AKras* mice. DAPI was used as counterstain. Scale bar = 200μm. P) Quantification of numbers of SOX2⁺ cells in the ventral DG of AKras*(c) and Pdx1Cre-AKras* mice. n=3 AKras*(c) mice and n=5 Pdx1Cre-AKras* mice. Data were analyzed with the two-tailed unpaired student’s t-test. Error bars represent SEM. Q) Representative images of SOX2^+^DCX^+^ cells in the ventral DG of AKras*(c) and Pdx1Cre-AKras* mice. Scale bar = 200μm. R) Quantification of numbers of SOX2^+^DCX⁺ cells in the ventral DG of AKras*(c) and Pdx1Cre-AKras* mice. n=3 AKras*(c) mice and n=5 Pdx1Cre-AKras* mice. Data were analyzed with the two-tailed unpaired student’s t-test. Error bars represent SEM.

Taking advantage of this well-defined temporal progression, we asked whether adult hippocampal neurogenesis is differentially affected during distinct stages of tumor development. At P21, corresponding to early tumor development, no significant differences were observed between Pdx1Cre-AKras* mice and control littermates in the number of DCX⁺ immature neurons or GFAP⁺ radial glia-like (RGL) neural stem cells, in either the dorsal or ventral dentate gyrus (DG) (Fig. S3E–J). These findings indicate preserved neurogenic activity and NSC pool size at early stages of disease.

In contrast, at P40, when tumors were more advanced, Pdx1Cre-AKras* mice exhibited a significant reduction in adult neurogenesis. In the dorsal dentate gyrus (DG), the number of DCX⁺ immature neurons was markedly decreased compared to control littermates (Fig. 3C,D). This was accompanied by a reduction in GFAP⁺ radial glia-like (RGL) neural stem cells, indicating a primary significant loss of hippocampal NSC (Fig. 3E,F). In contrast, the total number of SOX2⁺ cells was not altered (Fig. 3G,H), suggesting that the overall progenitor population may be partially preserved. Despite this, the number of SOX2⁺/DCX⁺ double-positive cells was significantly reduced (Fig. 3I,J), indicating impaired survival or/and neuronal differentiation potential along the neurogenic lineage. Analysis of the ventral DG revealed a similar phenotype, with reduced numbers of DCX⁺ cells and GFAP⁺ RGL NSCs (Fig. 3K-N). SOX2⁺ cell numbers remained largely unchanged, however the number of SOX2⁺/DCX⁺ double-positive cells was again significantly reduced (Fig. 3O-R). In line with this, the ratio of SOX2⁺/DCX⁺ cells relative to total SOX2⁺ cells showed a decreasing trend in both dorsal and ventral DG, although these differences did not reach statistical significance (Fig. S3K,L).

To exclude potential confounding effects of systemic metabolic alterations, we monitored body weight, blood glucose, and circulating insulin levels. While Pdx1Cre-AKras* mice displayed reduced body weight and altered glycemia at later stages, insulin levels were not significantly different between groups (Fig. S3B–D), suggesting that impaired neurogenesis is unlikely to be driven by insulin deficiency.

Moreover, to exclude potential off-target recombination, we crossed Pdx1^Cre^ mice with ROSA26^lsl-tdTomato^ reporter mice. In the brain, tdTomato expression was restricted to the hypothalamus, with no detectable signal in the hippocampus (Fig. S3A), ruling out ectopic Cre activity and oncogenic Kras expression in hippocampal cells.

Finally, Evans Blue extravasation assays did not reveal significant differences in blood–brain barrier (BBB) permeability between control and tumor-bearing mice (Fig. S3M).

Together, these findings demonstrate that adult hippocampal neurogenesis is preserved during early stages of pancreatic tumor development but becomes significantly impaired as the disease progresses. Notably, while both xenograft and genetic models exhibit reduced neurogenesis, the differential effects on SOX2⁺ cell populations suggest that distinct factors or/and mechanisms may contribute to NSC dysfunction across models.

### Systemic factors from tumor-bearing mice impair hippocampal NSC survival independent of major neuroimmune alterations

Crosstalk between the periphery and the brain is known to regulate adult hippocampal neurogenesis, in part through circulating factors such as cytokines. To determine whether systemic signals contribute to the observed neurogenic defects in pancreatic cancer (PC), we directly tested the impact of circulating factors on neural stem cell (NSC) function in vitro. Serum was collected from tumor-bearing mice in both models (Xeno^Panc1^ and Pdx1Cre-AKras*) and applied to primary hippocampal NSCs isolated from wild-type mice and cultured as neurospheres (Fig. 4A). After expansion, neurospheres were dissociated and plated under conditions either supporting proliferation (presence of EGF and FGF2) or inducing apoptosis (growth factor withdrawal).

**Fig 4.**
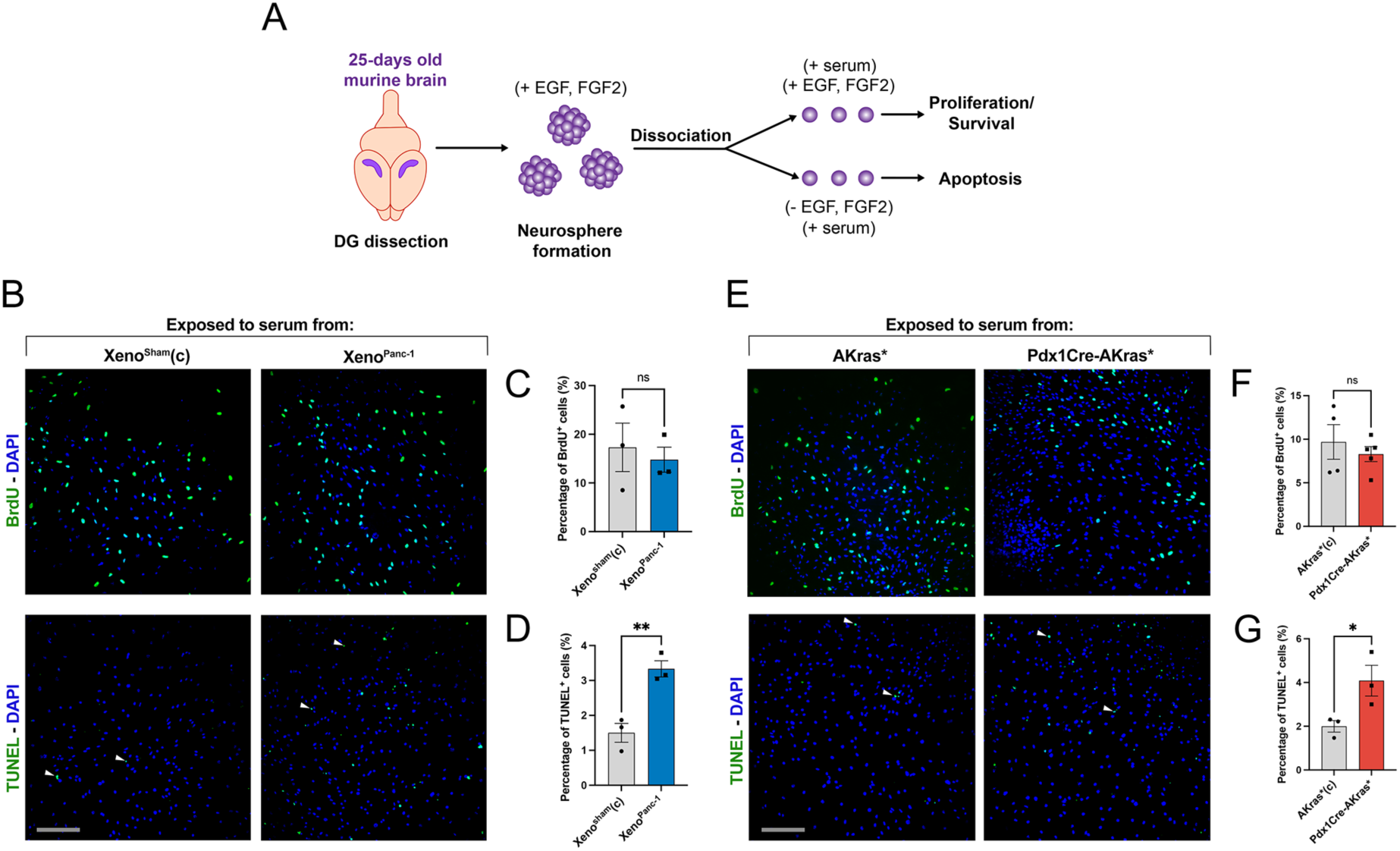
Systemic factors alter NSC behavior ex vivo, leading to reduced survival. A) Schematic representation of the experimental workflow for neurosphere culture. B) Representative images of wt C57BL/6 isolated NSCs exposed to serum from Xeno^Sham(c)^ and Xeno^Panc-1^ mice and stained for BrdU after 2-hour BrdU incorporation and assessed by TUNEL. DAPI was used as a counterstain. Scale bar = 200μm. C) Quantification of the numbers of BrdU^+^ cells exposed to serum from Xeno^Sham(c)^ and Xeno^Panc-1^ mice. n=3 biological replicates for each group. Data were analyzed with the two-tailed unpaired student’s t-test. Error bars represent SEM. D) Quantification of apoptotic cells as assessed by TUNEL. n=3 biological replicates for each group. Data were analyzed with the two-tailed unpaired student’s t-test. Error bars represent SEM. E) Representative images of wt C57BL/6 isolated NSCs exposed to serum from AKras*(c) and Pdx1Cre-AKras* mice and stained for BrdU after 2-hour BrdU incorporation and assessed by TUNEL. DAPI was used as a counterstain. Scale bar = 200μm. F) Quantification of the numbers of BrdU^+^ cells exposed to serum from AKras*(c) and Pdx1Cre-AKras* mice. n=4 replicates exposed to serum from AKras*(c) mice and n=5 replicates exposed to serum from Pdx1Cre-AKras* mice. Data were analyzed with the two-tailed unpaired student’s t-test. Error bars represent SEM. G) Quantification of apoptotic cells as assessed by TUNEL. n=3 biological replicates for each group. Data were analyzed with the two-tailed unpaired student’s t-test. Error bars represent SEM.

Exposure to serum from tumor-bearing mice did not significantly affect NSC proliferation, as assessed by BrdU incorporation, in either model (Fig. 4B, C, E, F). In contrast, NSC survival was markedly impaired, as indicated by a significant increase in TUNEL-positive cells following treatment with tumor-derived serum (Fig. 4B, D, E, G). These results demonstrate that circulating factors in PC selectively compromise NSC survival without affecting proliferative capacity.

To further assess whether local neuroimmune changes could contribute to impaired neurogenesis, we analyzed microglial abundance and activation in the hippocampus. Immunofluorescence staining for IBA1 revealed no major differences in microglial cell numbers in either the dentate gyrus (DG) or hilus across both models, with the exception of a modest increase in the dorsal DG of Xeno^Panc1^ mice (Fig. S4A–J). Consistent with this, qPCR analysis of hippocampal tissue from Pdx1Cre-AKras* mice showed no significant changes in the expression of Cd11b or MertK suggesting non-inflammatory homeostatic status of microglia, while Cd68 expression that is linked with activated states of microglia was modestly reduced.

Together, these findings demonstrate that circulating factors associated with pancreatic tumors selectively impair hippocampal NSC survival, while local neuroimmune alterations remain modest, suggesting that systemic rather than local inflammatory mechanisms underlie the observed deficits in adult hippocampal neurogenesis.

### Distinct systemic inflammatory profiles across PC models converge on IL-6 as a candidate mediator of impaired AN

To identify circulating factors that may underlie the observed impairment in adult hippocampal neurogenesis, we performed multiplex cytokine profiling using a Luminex-based panel measuring 20 inflammatory mediators in serum from both pancreatic cancer (PC) models. In the genetic model, serum was collected from Pdx1Cre-AKras* mice at P21 and P40, representing early and advanced disease stages, respectively. In parallel, sera from Xeno^Panc1^ mice were collected at developing and advanced tumor stages.

Comparative analysis revealed that the systemic inflammatory profiles differed markedly between the two models (Fig. 5A). In Xeno^Panc1^ mice, G-CSF, MCP-1, and RANK were elevated during early tumor development, whereas in advanced disease these factors, together with IL-6, were among the most prominently upregulated cytokines (Fig. 5A,B). In contrast, the genetic Pdx1Cre-AKras* model displayed a broader and stage-dependent inflammatory response. At P21, a limited set of cytokines, including IL-17F, M-CSF, TNFSF11, IL-1α, and IL-12p40, were increased. By P40, most measured cytokines were elevated, with TNFα, IL-1α, IL-1β, and IL-12p40 showing the strongest upregulation (Fig. 5A,C). Notably, although IL-6 was not among the most abundantly expressed cytokines in this model, its circulating levels were substantially elevated at P40 compared to controls (Fig. 5C). Importantly, IL-6 emerged as the only cytokine consistently elevated across both pancreatic cancer models, highlighting it as a potential common systemic mediator associated with tumor progression.

**Fig 5.**
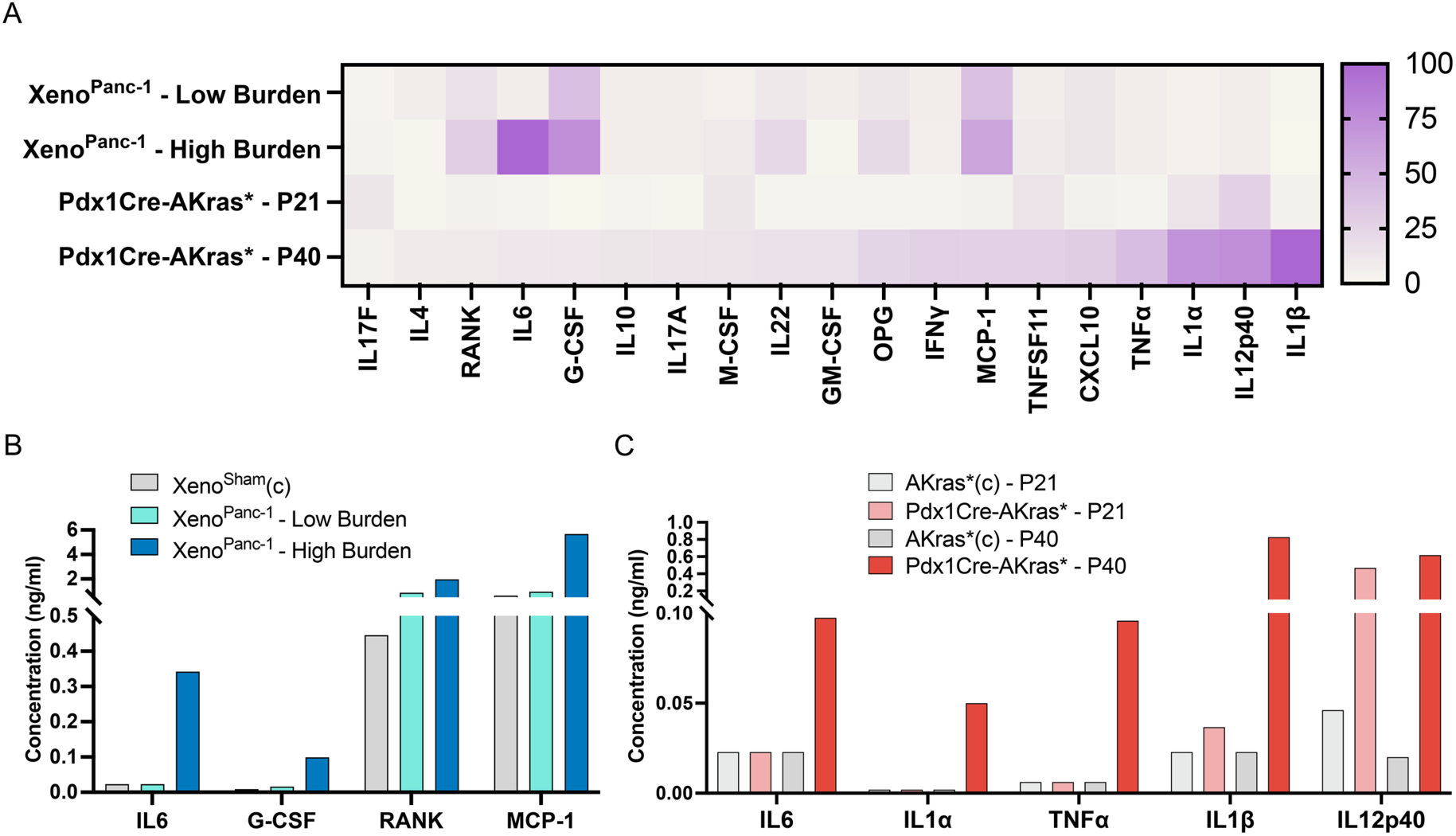
Proinflammatory cytokines are upregulated in mice with PC, with distinct profiles between the two mouse models. A) Heatmap displaying normalized fold changes of the concentration of selected cytokines in the serum of Xeno^Panc-1^ – Low Burden against Xeno^Sham(c)^, Xeno^Panc-1^ – High Burden against Xeno^Sham(c)^, Pdx1Cre-AKras* - P21 against AKras*(c) - P21 and Pdx1Cre-AKras* - P40 against AKras*(c) - P40. B) Quantification of the concentration of IL-6, MCP-1, G-CSF, and RANK in the serum of Xeno^Sham(c)^, Xeno^Panc-1^ – Low Burden and Xeno^Panc-1^ – High Burden mice. C) Quantification of the concentration of IL6, IL1α, TNFα, IL1β, and IL12p40 in the serum of AKras*(c) - P21, Pdx1Cre-AKras* - P21, AKras*(c) - P40 and Pdx1Cre-AKras* - P40 mice.

To determine whether these systemic changes are reflected locally within the hippocampus, we analyzed the expression of selected inflammatory genes in hippocampal tissue from Pdx1Cre-AKras* mice. Surprisingly, Il6 mRNA levels were not altered, suggesting that elevated circulating IL-6 does not correspond to increased hippocampal local production but rather to a systemically increased production (Fig. S5A). Moreover, Il1a expression was reduced, while Il1b, Tnfa, Il12p40, Cxcl10, and Opg levels remained unchanged (Fig. S5B–G). These results indicate that systemic inflammatory signals are not broadly recapitulated at the transcriptional level within the hippocampus.

Collectively, these data support a model in which pancreatic cancer induces distinct systemic inflammatory environments that are not mirrored by overt local neuroinflammation, reinforcing the notion that peripheral cytokines act as key mediators of impaired adult hippocampal neurogenesis. Notably, despite clear differences in the broader cytokine profiles, potentially reflecting the distinct immune contexts of the two models, IL-6 emerged as a consistently elevated factor across both settings, positioning it as a particularly compelling candidate linking systemic inflammation to neurogenic deficits in pancreatic cancer.

## DISCUSSION

Pancreatic cancer (PC) carries one of the highest rates of depression among all malignancies, with psychiatric symptoms often preceding diagnosis by months or years ^1^. Despite this long-recognized clinical association, the biological mechanisms linking pancreatic tumor development to CNS dysfunction remain poorly understood. Here, using complementary xenograft and genetically engineered mouse models, we demonstrate that pancreatic tumor development drives a depressive-like behavioral phenotype and impairs adult hippocampal neurogenesis. By integrating cytokine profiling with functional in vitro assays, we identify systemic inflammatory signals, including IL-6, as candidate mediators of this process. Notably, depressive-like behavior emerged during early stages of disease in the xenograft model, mirroring the clinical observation that neuropsychiatric symptoms can precede pancreatic cancer diagnosis and suggesting that tumor-derived signals affect brain function remarkably early during disease progression.

The behavioral phenotype observed in XenoPanc1 mice strongly indicates depression-related behavior and is accompanied by a significant reduction in adult hippocampal neurogenesis. According to the neurogenic hypothesis of depression, adult-born neurons contribute to mood regulation and stress resilience, whereas impaired neurogenesis has been linked to depressive states and reduced antidepressant efficacy ^20^. Consistent with this framework, tumor-bearing mice exhibited reduced numbers of immature neurons and neural stem/progenitor cells in both dorsal and ventral dentate gyrus. While the ventral hippocampus is more directly linked to emotional behavior and HPA-axis regulation, the dorsal region is primarily associated with learning and memory ^31^. The broad neurogenic impairment observed across both regions therefore provides a plausible cellular substrate for the depressive-like phenotype. Furthermore, the absence of major changes in the SOX2+/DCX+ ratio suggests that the primary defect may involve depletion and survival of the neurogenic pool rather than a specific block in neuronal differentiation.

Validation in the Pdx1Cre-AKras* model strengthens these findings considerably. Neurogenesis was preserved during early disease stages but became significantly impaired as tumors progressed, indicating a temporal relationship between tumor burden and neurogenic dysfunction. Although some differences were observed between the xenograft and genetic models, particularly regarding SOX2+ populations, both converged on the same principal outcome: reduced neurogenic activity and depletion of the neural stem cell compartment. These differences may reflect the distinct immune environments of immunodeficient and immunocompetent mice or differences in the composition of systemic inflammatory signals. Importantly, several alternative explanations were excluded. Comparable insulin levels argue against insulin deficiency as a major contributor, blood-brain barrier integrity remained intact, and ectopic hippocampal Kras activation was excluded in the genetic model. Together, these findings support the conclusion that pancreatic tumors influence the neurogenic niche through indirect systemic mechanisms.

An important mechanistic insight emerged from the serum-transfer experiments. Exposure of primary hippocampal NSCs to serum derived from tumor-bearing mice selectively increased apoptosis without significantly affecting proliferation, indicating a pro-survival rather than anti-proliferative defect. These findings are consistent with the in vivo observations and place our work within an emerging paradigm in which adult NSCs act as sensors of systemic physiological state and respond dynamically to circulating factors generated during disease and inflammation. Similar effects on adult neurogenesis have been reported in other pathological conditions, including COVID-19-associated neurological disease, systemic autoimmune disorders and chronic inflammatory diseases ^32–34^.

Among the inflammatory mediators profiled, IL-6 emerged as the only cytokine consistently elevated across both models at disease stages associated with neurogenic impairment. This is notable given the established links between IL-6, pancreatic cancer progression and depression, as well as evidence that IL-6 negatively regulates adult neurogenesis ^35^. Moreover, our recent work in neuropsychiatric lupus demonstrated that circulating IL-6 can directly induce apoptosis of adult hippocampal NSCs through a BBB-independent mechanism, providing a compelling mechanistic precedent for the present findings ^33^. Although our data do not establish a causal role for IL-6, they support the hypothesis that systemic IL-6 contributes to communication between pancreatic tumors and the hippocampal neurogenic niche. Importantly, elevated circulating cytokines were not mirrored by widespread inflammatory gene expression within the hippocampus, suggesting that peripheral rather than local inflammatory mechanisms predominate. Other cytokines elevated in the genetic model, including TNFα, IL-1α and IL-1β, may also contribute directly or indirectly to NSC dysfunction, indicating that pancreatic cancer-associated neurogenic impairment is likely driven by a broader inflammatory network rather than a single cytokine.

To address why this phenomenon occurs in PC, it is important to consider the systemic consequences of tumor progression. Pancreatic cancer behaves as a highly inflammatory disease, particularly at advanced stages, resulting in the release of numerous cytokines and immune mediators into the circulation. Our findings suggest that NSCs may be particularly sensitive to these systemic changes and that chronic exposure to inflammatory signals compromises the maintenance of the neurogenic niche. Whether this represents an adaptive host response or a consequence of tumor-driven systemic inflammation remains unknown.

Taken together, our findings support a model in which pancreatic tumor development drives depressive-like behavior through systemic inflammatory signals that impair hippocampal NSC survival and disrupt adult neurogenesis. These observations position PCAD as a biological consequence of tumor progression rather than merely a psychological response to diagnosis and suggest that preserving neurogenesis may represent a novel therapeutic avenue. While IL-6-targeted therapies represent one potential strategy, broader approaches aimed at reducing systemic inflammation or promoting neurogenesis - including antidepressants, physical exercise, environmental enrichment, and interventions targeting neuroimmune signaling pathways - may also prove beneficial.

Beyond improving mood and quality of life, alleviating depression may have direct consequences for cancer progression itself. Increasing evidence supports extensive bidirectional communication between the brain and immune system, whereby chronic stress and depressive states impair anti-tumor immunity through effects on cytotoxic T cells, natural killer cells, and inflammatory signaling networks ^36,37^. Conversely, restoration of normal hippocampal function and adult neurogenesis has been associated with improved immune regulation and resilience to inflammatory challenges ^38^. Although the relationship between adult neurogenesis and tumor progression remains unexplored in pancreatic cancer, our findings raise the intriguing possibility that disruption of the hippocampal neurogenic niche may represent more than a neuropsychiatric consequence of disease. Specifically, pancreatic tumors may actively suppress adult hippocampal neurogenesis as part of a systemic adaptation that ultimately favors immune evasion. Conversely, preservation of the neurogenic niche may contribute not only to improved psychological well-being but also to enhanced immune-mediated tumor control. Future studies should therefore investigate whether interventions that preserve or restore adult hippocampal neurogenesis can simultaneously alleviate pancreatic cancer-associated depression and improve anti-tumor immune responses.

## METHODS

### Mouse strains, maintenance and genotyping

Animal maintenance and experimentation were conducted in accordance with the FELASA recommendations and the ethical and practical guidelines for the care and use of laboratory animals set by the competent veterinary authorities in the authors’ institutions. Experiments were all conducted in both male and female mice and no differences were observed.

Preexisting mouse strains and transgenic mice used were *Gt(ROSA)26Sor^tm9(CAG-tdTomato)hZE^ (ROSA26^lsl-tdTomato^,* reporter line that expresse*s tdTomato* under Cre Recombination*), Tg^(Pdx1-cre)6Tuv/J^ (Pdx1^Cre^,* expresses Cre Recombinase under *Pdx1* promoter) and Actb^tm1(KrasG12D)Arge^ (AKRas*, expresses the oncogenic *KRAS4B^G12D^* placed in the *Actb* locus) ^30,39,40^.

Genotyping was performed by conventional PCR on genomic DNA isolated from mouse-tails using standard procedures. Briefly, mouse tails were dissolved in Tail buffer (100 mM Tris-HCl pH 8.5, 200 mM NaCl, 5 mM EDTA and 0.2% SDS) with 50 μg/ml Proteinase K (Sigma) overnight at 55°C. Following a protein extraction step with phenol/chloroform (Sigma), genomic DNA was precipitated from the aqueous phase with 100% ethanol and finally suspended in TE buffer (10 mM Tris-HCI pH 8.5 and 1 mM EDTA). PCR primers for mouse genotyping are provided in Supplementary Table S1.

### Orthotopic xenografts of pancreatic cancer

Xenografts were generated in NOD-SCID (Xeno^Panc1^) mice through implantation of the human pancreatic cancer cell line Panc-1 (1.5 × 10⁵ cells suspended in 40 μl of serum-free medium) ^41^, representing the Xeno^Panc1^ - High Burden cohort of mice. Following a midline incision of the anterior abdominal wall, the cell suspension was directly injected into the pancreatic parenchyma. Most animals were euthanized 14 days post-injection (p.i), with no mortality observed. For preliminary tumor assessment, subsets of mice were sacrificed at 5, 11, and 21 days p.i. An equal number of sham-operated mice was included as a control group.

In a separate experiment, a second cohort of mice representing earlier stages of tumor development, termed Xeno^Panc1^ - Low Burden, was established. The procedure was identical except for the initial cell inoculum, which was reduced to 5 × 10⁴ cells in 40 μl of serum-free medium.

### Behavioral experiments

Behavioral experiments were conducted 11-13 days post-surgery for both tumor-bearing and sham-operated NOD-SCID mice. At this stage, the tumor was not palpable and the mice did not exhibit any kind of distress. For the Open-Field Test, mice were placed into the Open Field arena for a total of 5 minutes. Their movement was video-recorded and analyzed by the software (EthoVisionXT) calculating the total distance travelled and the time spent in the center zone of the arena. For the Elevated Plus Maze Test, mice were placed at the center of the maze where they faced a closed arm at the beginning of the test and the testing lasted a total of 5 minutes. Their movement was video-recorded and analyzed by the software (EthoVisionXT) calculating the total distance travelled and the time spent in the open arms of the Maze. For the Tail Suspension Test, mice were carefully affixed by their tail using laboratory tape to a horizontal shelf and their testing lasted for a total of 6 minutes. Their movement was video-recorded and analyzed by the software (EthoVisionXT) calculating their immobility time.

### Immunostainings on cryosection and histological analysis

For immunofluorescence (IF), sections were post-fixed for 10 min with 4% w/v PFA and then washed in PBS (3 washes, 5 min each). Heat-Induced Antigen Retrieval was performed using 10 mM Sodium Citrate (pH 6), for 15 min in a microwave at nearly boiling temperature. Sections were allowed to cool down to room temperature (RT), washed in PBS (3 washes, 5 min each) and incubated overnight (O/N) at 4 °C with the primary antibodies diluted in 10% w/v NGS, 0.3% v/v Triton-X in PBS. The next day, sections were washed in PBS (3 washes, 5 min each) and incubated for 1hr at RT with the secondary antibodies diluted in 10% w/v NGS, 0.3% v/v Triton-X in PBS with DAPI (Invitrogen, 1:1000). Sections were washed in PBS (3 washes, 5 min each) and mounted with Fluoroshield (Sigma).

Cells were washed three times with PBS incubated in blocking buffer (10% w/v NGS, 0,1% v/v Triton-X in PBS) for 1hr at RT. For BrdU staining, the blocking step was replaced with incubation with HCl 2N at 37 °C for 15 min followed by 20 min incubation with Sodium Borate (pH=8.5) at RT. Primary antibodies were diluted in 10% w/v NGS, 0,1% v/v Triton-X in PBS and incubated O/N at 4 °C. Cells were washed three times with PBS for 5 min and incubated with the secondary antibodies diluted in 10% w/v NGS, 0,1% v/v Triton-X in PBS for 1hr. Following three washes with PBS for 5 min, cells were incubated with DAPI (1:1000) diluted in 10% w/v NGS, 0,1% v/v Triton-X in PBS for 10 min, washed with PBS (3 times, 5 min each) and finally mounted onto slides using Fluoroshield. All the antibodies used are listed in Supplementary Table S2.

For immunohistochemical analysis, paraffin-embedded pancreatic tissues were sectioned, deparaffinized/rehydrated, stained for hematoxylin and eosin, dehydrated and mounted following standard procedures.

### TUNEL assay

Cells were permeabilized with 0.2% Triton-X100 in PBS for 5 min. After a PBS wash, coverslips were incubated in equilibration buffer for 10 min. The mix with the nucleotides and the rTdT enzyme was then added to the coverslips for 1 hr at 37°C. Following inactivation with SSC 2x for 10 min at RT, cells were washed in PBS, incubated with DAPI (1:1000 dilluted in PBS), washed again in PBS and finally mounted onto slides using Fluoroshield.

### Microscopy

Images were acquired with a Leica TCS SP5 inverted confocal (20×0.7NA dry objective lens) and captured using camera software (LAS AF, Leica Microsystems). Images were acquired as z-stacks with a z-step of 1 μm.

### Morphometric analysis of immunofluorescent images

Images were processed in ImageJ version 2.0.0-rc-59/1.51n (imagej.nih.gov). Cell counting was performed manually. Image panels were assembled in Photoshop 2023 24.3.0 Release (Adobe).

### Mouse Adult Hippocampal NSCs

For the ex vivo experiment, 25 days old C57BL/6 mice were euthanized and their brains were removed. The DG was micro-dissected from the rest of the hippocampus, digested with a Trypsin-based solution and mechanically triturated. Medium was renewed every 4 days and the culture was maintained for 14-18 days in order to develop neurospheres. When they reached the desired size, floating neurospheres were trypsin-dissociated, triturated into single-cells, plated onto poly-L-lysine coated coverslips in 24-well plates and cultured in the presence (proliferation assay) or absence (survival assay) of EGF/bFGF. Isolated sera from tumor-bearing and control mice were added to the culture one day after plating and maintained for 24 h. Cells were then fixed for 10 minutes in PFA 4% and stored in PBS 1x until immunofluorescence. For the proliferation study BrdU (10 μM) was added 2h before fixation.

### Serum isolation

Mice were anesthetized at the desired age and blood was collected through cardiac puncture. Whole blood was allowed to clot by standing for 30 minutes at room temperature and centrifuged at 4°C/2000rpm/30min to pellet cells.

### Cytokine and hormone detection

A Luminex-based multiplex panel was used to quantify cytokine levels in the isolated sera. Measured cytokines were OPG, IL-17F, IL-6, GM-CSF, IFN-g, IL-10, IL-1a, IL-4, CXCL10, MCP-1, TNFa, M-CSF, G-CSF, IL-22, IL-1b, TNFSF11, RANK, IL-17A and IL-12p40. A total of 12 samples was used (2 samples of Pdx1Cre-AKRas^G12D^ mice at P40, 2 samples of AKRas^G12D^mice at P40, 1 sample of Pdx1Cre-AKRas^G12D^ mice at P21, 1 sample of AKRas^G12D^ mice at P21, 2 samples of Xeno^Panc1^ - High Burden, 2 samples of Xeno^Panc1^ - Low Burden, 2 samples of sham-operated NOD-SCID controls). To detect Insulin levels, sera were assayed using the appropriate ELISA kit (Mercodia; 10-1249) and following the manufacturer’s instructions.

### Blood glucose measurements

Pdx1Cre-AKRas^G12D^ mice and their respective controls were used to measure their blood glucose levels following a 16-hour overnight fasting. Animals were weighed and glucose concentration was measured from a tail-vein blood drop using a handheld glucometer (Contour Next; Bayer) according to the manufacturer’s instructions.

### BBB permeability assay

Blood–brain barrier (BBB) permeability was assessed with Blue Evans assay (EB). Pdx1Cre-AKRas^G12D^ mice and their respective controls received an intravenous tail injection of EB (25 mg/kg in PBS) at P40. After After 4 hours, the hippocampus was isolated from each animal, while lung tissue from two mice per group served as a positive control. Collected tissues were incubated overnight at 60°C in formamide (5 mL per mg of tissue) to extract the dye. EB levels were quantified using an ELISA reader at 620 nm. Concentrations were calculated with the help of a calibration curve generated from the absorbance values of seven samples with known Evans Blue concentrations, as previously described ^33^.

### RNA extraction, cDNA synthesis and qRT-PCR

Hippocampal tissue was dissected from Pdx1Cre-AKRas^G12D^ mice and their respective controls. Total RNA was extracted using TRI Reagent following a standard protocol. Briefly, tissue samples were lysed in TRI Reagent and homogenized thoroughly. Chloroform was added, followed by centrifugation to achieve phase separation. The aqueous (upper) phase was collected and mixed with isopropanol, then incubated overnight at −80°C to precipitate RNA. After centrifugation, the supernatant was discarded and the RNA pellet was washed with ethanol and centrifuged again. The pellet was air-dried and finally resuspended in DEPC-treated water. Reactions were carried out from four independent samples. Absolute expression values were calculated using the ΔCt method using *β-actin* for normalization. Primers that were used for real-time qPCR are listed in Supplementary Table S3.

### Statistical analysis

Data are expressed as means and presented in bar graphs, line charts and heatmaps generated using the GraphPad Prism Software (v. 10). Error bars represent Standard Error of Mean (SEM). Normality of the data distribution was assessed using the Shapiro-Wilk test. Statistical significance for two-sample comparisons was determined by two-tailed, unpaired Student’s t tests (parametric testing), or by Wilcoxon rank-sum/Mann-Whitney U tests (non-parametric testing). For comparisons involving more than two groups, statistical significance was determined using two-way ANOVA. Differences were considered statistically significant if the p-value was less than 0.05 (*p < 0.05, **p < 0.01, ***p < 0.005). The sample size (n) for each experiment is indicated in the corresponding figure legend and, unless otherwise stated, it refers to biological replicates.

## AUTHOR CONTRIBUTIONS

N.N., Y.K. and I.S. conceived the study and Y.K. and I.S. designed the experiments. D.T., A.P., C.C. and P.K. performed the investigation. D.T., A.P., A.P.(2), Y.K. and I.S. conducted the analysis and interpretation of results. D.T. wrote the original draft, with critical revision from Y.K. and I.S. All authors reviewed, edited and approved the final manuscript. Y.K. and I.S. take primary responsibility for the data described in this manuscript.

## ACKNOWLDEGMENTS

This work was financially supported by the Empirikion Foundation (CRA No. 2024/14 to I.S.), and a Fondation Sante Sidney Altman Scholarship awarded to D.T. We thank Argiris Efstratiadis (BRFAA) for his generous gift of the Actb^LSLKras*^ mouse line. We thank Pavlos Alexakos and Vagelis Balafas of the BRFAA Animal Facility for expert animal husbandry and invaluable assistance with mouse handling procedures. We are also grateful to Effrosyni Koronaiou for her guidance in establishing the behavioral assays. Finally, we acknowledge the support of the BRFAA Bioimaging Unit, particularly Stamatis Pagkakis and Anastasios Dellis for their technical assistance, and the BRFAA Histology Unit, especially Anna Agapaki, for expert assistance with paraffin tissue processing.

