## Supplemental Figures S1-S5 and Tables 1-3 for "Pancreatic cancer disrupts the adult hippocampal neurogenic niche"

### SUPPLEMENTARY FIGURES AND TABLES

***Figure S1***

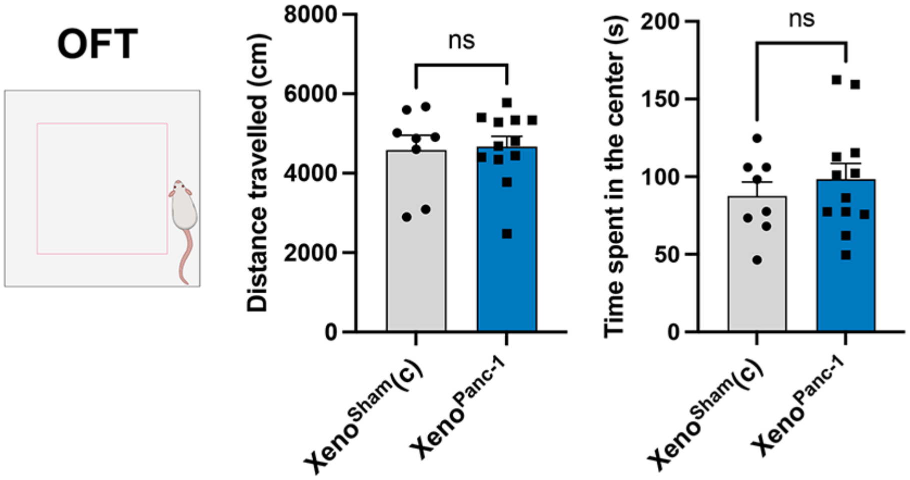


**Fig S1. Motility of Xeno$^{Panc\text{-}1}$ mice is not affected by PC.**

Quantification of mobility (distance travelled in the arena) and anxiety-phenotype (time spent in center zone) in the Open Field Test. n=8 Xeno$^{Sham(c)}$ mice and n=10 Xeno$^{Panc\text{-}1}$ mice. Data were analyzed with the two-tailed unpaired student’s t-test. Error bars represent SEM.

*Figure S2*

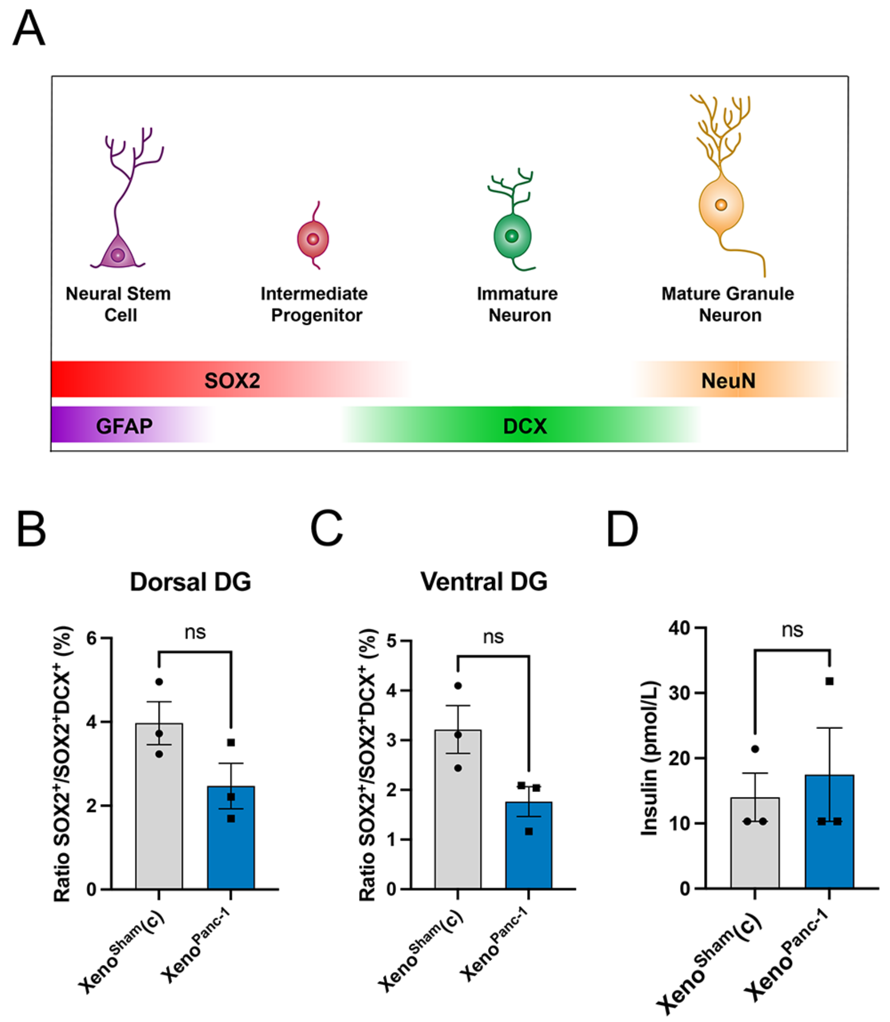


**Figure S2. The differentiation rate of SOX2⁺ proliferating cells is not significantly altered in Xeno$^{Panc-1}$ mice.**

A) Schematic representation of AN stages and their associated cell markers. B) Quantification of the ratio of $SOX2^+$ to $SOX2^+DCX^+$ cells in the dorsal DG of Xeno$^{Sham(c)}$ and Xeno$^{Panc-1}$ mice. n=3 for each group. Data were analyzed with the two-tailed unpaired student's t-test. Error bars represent SEM. C) Quantification of the ratio of $SOX2^+$ to $SOX2^+DCX^+$ cells in the ventral DG of Xeno$^{Sham(c)}$ and Xeno$^{Panc-1}$ mice. n=3 for each group. Data were analyzed with the two-tailed unpaired student's t-test. Error bars represent SEM. D) Quantification of Insulin concentration levels in the serum of Xeno$^{Sham(c)}$ and Xeno$^{Panc-1}$ mice. n=3 for each group. Data were analyzed with the Mann Whitney test. Error bars represent SEM.

*Figure S3*

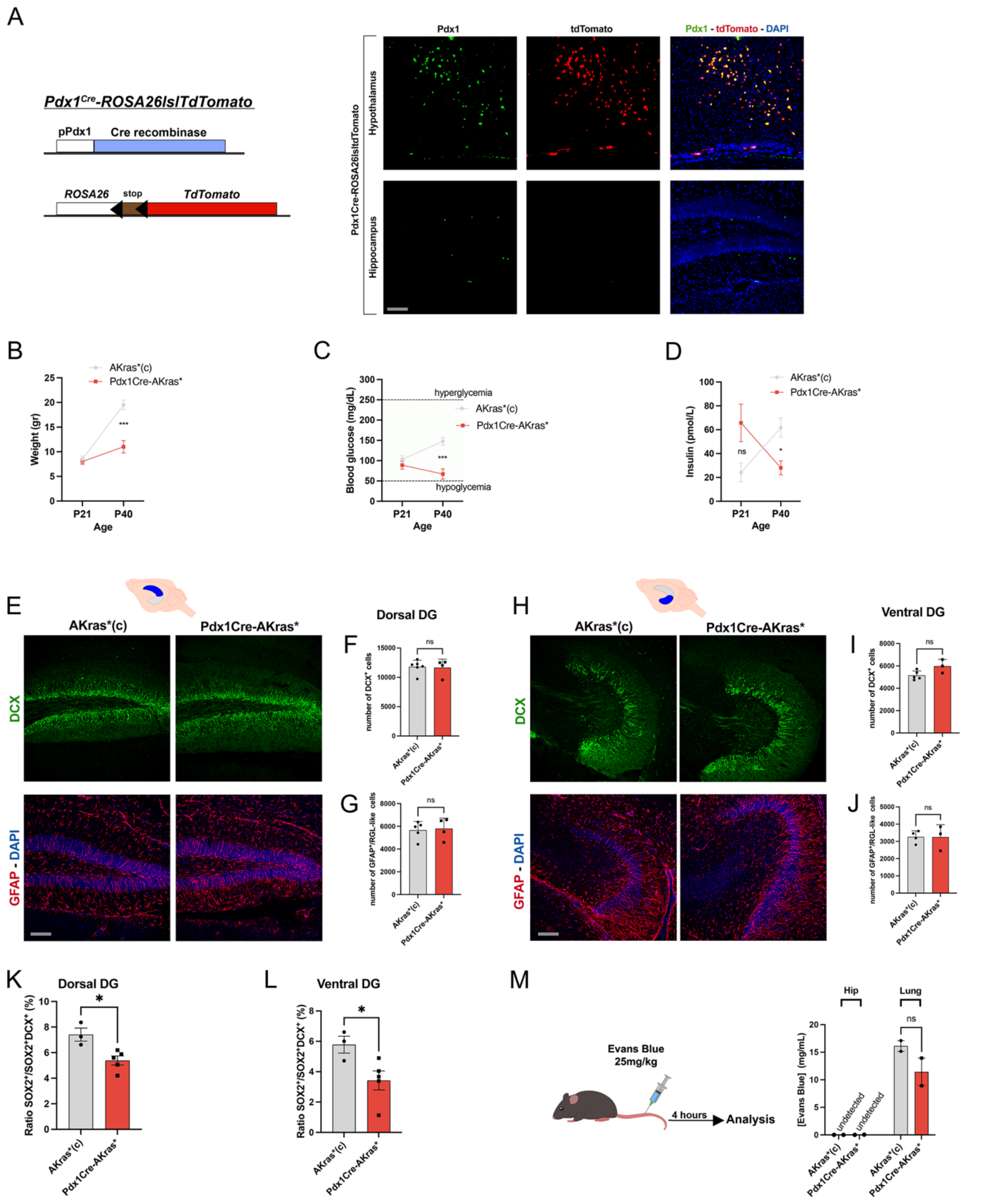


**Fig S3. Pdx1Cre-AKras* mice exhibit normal metabolism and neurogenic capacity at early postnatal stages.**

A) Schematic representation of Pdx1Cre-Rosa26lsltdTomato genotype and representative images of Pdx1$^{+}$ and tdTomato$^{+}$ cells in the Hypothalamus and Hippocampus. DAPI was used as a counterstain. Scale bar = 200µm. B) Quantification of body weight of AKras*(c) and

Pdx1Cre-AKras* mice at P21 and P40. n=8 AKras*(c) mice and n=6 Pdx1Cre-AKras* mice. Data were analyzed with the two-tailed paired t-test. Error bars represent SEM. C) Quantification of blood glucose concentration of AKras*(c) and Pdx1Cre-AKras* mice at P21 and P40. n=8 AKras*(c) mice and n=6 Pdx1Cre-AKras* mice. Data were analyzed with two-way ANOVA. Error bars represent SEM. D) Quantification of Insulin concentration levels in the serum of AKras*(c) and Pdx1Cre-AKras* mice at P21 and P40. n=4 for each group. Data were analyzed with two-way ANOVA. Error bars represent SEM. E) Representative images of $DCX^+$ cells and $GFAP^+$ cells in the dorsal DG of AKras*(c) and Pdx1Cre-AKras* mice at P21. DAPI was used as a counterstain. Scale bar = 200μm. F) Quantification of numbers of $DCX^+$ cells in the dorsal DG of AKras*(c) and Pdx1Cre-AKras* mice. n=6 AKras*(c) mice and n=4 Pdx1Cre-AKras* mice. Data were analyzed with the two-tailed unpaired student's t-test. Error bars represent SEM. G) Quantification of numbers of $GFAP^+$ cells in the dorsal DG of AKras*(c) and Pdx1Cre-AKras* mice. n=5 AKras*(c) mice and n=4 Pdx1Cre-AKras* mice. Data were analyzed with the two-tailed unpaired student's t-test. Error bars represent SEM. H) Representative images of $DCX^+$ cells and $GFAP^+$ cells in the ventral DG of AKras*(c) and Pdx1Cre-AKras* mice at P21. DAPI was used as a counterstain. Scale bar = 200μm. I) Quantification of numbers of $DCX^+$ cells in the ventral DG of AKras*(c) and Pdx1Cre-AKras* mice. n=5 AKras*(c) mice and n=3 Pdx1Cre-AKras* mice. Data were analyzed with the two-tailed unpaired student's t-test. Error bars represent SEM. J) Quantification of numbers of $GFAP^+$ cells in the ventral DG of AKras*(c) and Pdx1Cre-AKras* mice. n=4 AKras*(c) mice and n=3 Pdx1Cre-AKras* mice. Data were analyzed with the two-tailed unpaired student's t-test. Error bars represent SEM. K) Quantification of the ratio of $SOX2^+$ to $SOX2^+DCX^+$ cells in the dorsal DG of AKras*(c) and Pdx1Cre-AKras* mice. n=3 AKras*(c) mice and n=5 Pdx1Cre-AKras* mice. Data were analyzed with the two-tailed unpaired student's t-test. Error bars represent SEM. L) Quantification of the ratio of $SOX2^+$ to $SOX2^+DCX^+$ cells in the ventral DG of AKras*(c) and Pdx1Cre-AKras* mice. n=3 AKras*(c) mice and n=5 Pdx1Cre-AKras* mice. Data were analyzed with the two-tailed unpaired student's t-test. Error bars represent SEM. M) Quantification of EB in the Hippocampal and Lung tissue of AKras*(c) and Pdx1Cre-AKras* mice at P40. n=2 for each group. Data were analyzed with the Mann-Whitney test. Error bars represent SEM.

***Figure S4***

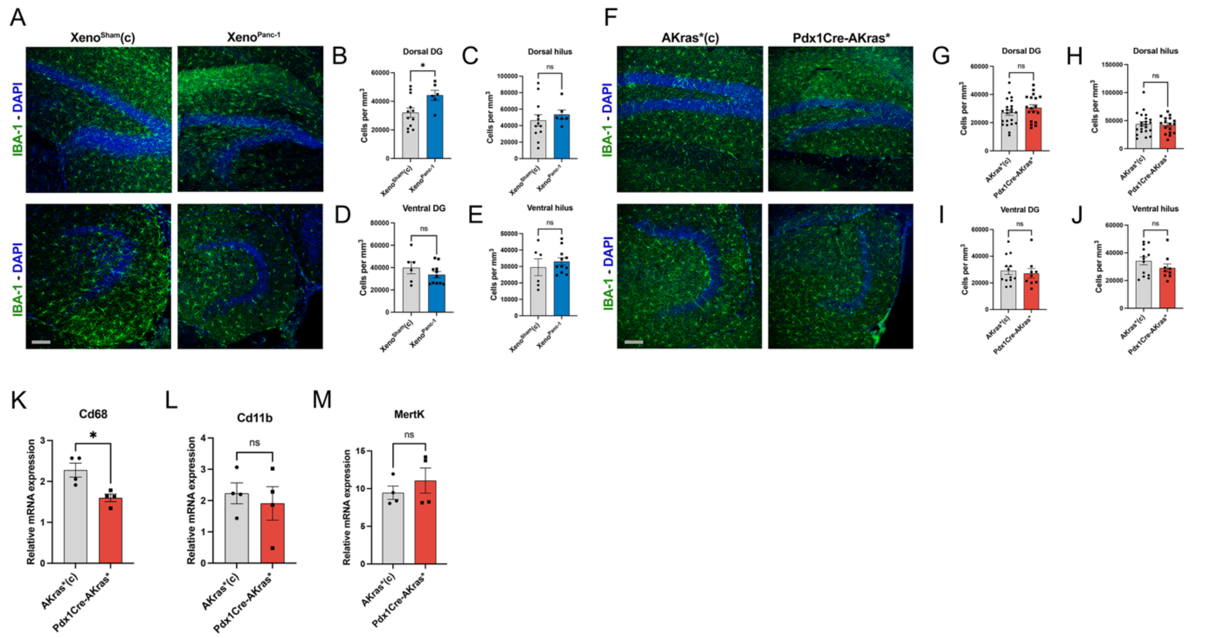


**Fig S4. Neuroinflammation is largely unchanged in mice with PC.**

**A)** Representative images of IBA1$^{+}$ cells in the dorsal and ventral DG of Xeno$^{Sham(c)}$ and Xeno$^{Panc\text{-}1}$ mice. DAPI was used as a counterstain. Scale bar = 200µm. **B)** Quantification of cell density of IBA1$^{+}$ cells in the dorsal DG of Xeno$^{Sham(c)}$ and Xeno$^{Panc\text{-}1}$ mice. n=12 sections from Xeno$^{Sham(c)}$ mice and n=6 sections from Xeno$^{Panc\text{-}1}$ mice. Data were analyzed with the two-tailed unpaired student's t-test. Error bars represent SEM. **C)** Quantification of cell density of IBA1$^{+}$ cells in the dorsal hilus of Xeno$^{Sham(c)}$ and Xeno$^{Panc\text{-}1}$ mice. n=12 sections from Xeno$^{Sham(c)}$ mice and n=6 sections from Xeno$^{Panc\text{-}1}$ mice. Data were analyzed with the two-tailed unpaired student's t-test. Error bars represent SEM. **D)** Quantification of cell density of IBA1$^{+}$ cells in the ventral DG of Xeno$^{Sham(c)}$ and Xeno$^{Panc\text{-}1}$ mice. n=6 sections from Xeno$^{Sham(c)}$ mice and n=11 sections from Xeno$^{Panc\text{-}1}$ mice. Data were analyzed with the two-tailed unpaired student's t-test. Error bars represent SEM. **E)** Quantification of cell density of IBA1$^{+}$ cells in the ventral hilus of Xeno$^{Sham(c)}$ and Xeno$^{Panc\text{-}1}$ mice. n=6 sections from Xeno$^{Sham(c)}$ mice and n=11 sections from Xeno$^{Panc\text{-}1}$ mice. Data were analyzed with the two-tailed unpaired student's t-test. Error bars represent SEM. **F)** Representative images of IBA1$^{+}$ cells in the dorsal and ventral DG of AKras*(c) and Pdx1Cre-AKras* mice. DAPI was used as a counterstain. Scale bar = 200µm. **G)** Quantification of cell density of IBA1$^{+}$ cells in the dorsal DG of AKras*(c) and Pdx1Cre-AKras* mice. n=22 sections from AKras*(c) mice and n=18 sections from Pdx1Cre-AKras* mice. Data were analyzed with the two-tailed unpaired student's t-test. Error bars represent SEM. **H)** Quantification of cell density of IBA1$^{+}$ cells in the dorsal hilus of AKras*(c) and Pdx1Cre-AKras* mice. n=22 sections from AKras*(c) mice and n=17 sections from Pdx1Cre-AKras* mice. Data were analyzed with the two-tailed unpaired student's t-test. Error bars represent SEM. **I)** Quantification of cell density of $IBA1^+$ cells in the ventral DG of AKras*(c) and Pdx1Cre-AKras* mice. n=13 sections from AKras*(c) mice and n=9 sections from Pdx1Cre-AKras* mice. Data were analyzed with the two-tailed unpaired student's t-test. Error bars represent SEM. **J)** Quantification of cell density of $IBA1^+$ cells in the ventral hilus of AKras*(c) and Pdx1Cre-AKras* mice. n=13 sections from AKras*(c) mice and n=9 sections from Pdx1Cre-AKras* mice. Data were analyzed with the two-tailed unpaired student's t-test. Error bars represent SEM. **K)** RT-qPCR analysis of expression levels of Cd68. Y-axis data is the relative mRNA expression ratio of AKras*(c) and Pdx1Cre-AKras* mice, and data of Pdx1Cre-AKras* mice is standardized to 1. P values were determined using two-tailed unpaired student's t-test. n=4 for each group. **L)** RT-qPCR analysis of expression levels of Cd11b. Y-axis data is the relative mRNA expression ratio of AKras*(c) and Pdx1Cre-AKras* mice, and data of Pdx1Cre-AKras* mice is standardized to 1. P values were determined using two-tailed unpaired student's t-test. n=4 for each group. **M)** RT-qPCR analysis of expression levels of MertK. Y-axis data is the relative mRNA expression ratio of AKras*(c) and Pdx1Cre-AKras* mice, and data of Pdx1Cre-AKras* mice is standardized to 1. P values were determined using two-tailed unpaired student's t-test. n=4 for each group.

*Figure S5*

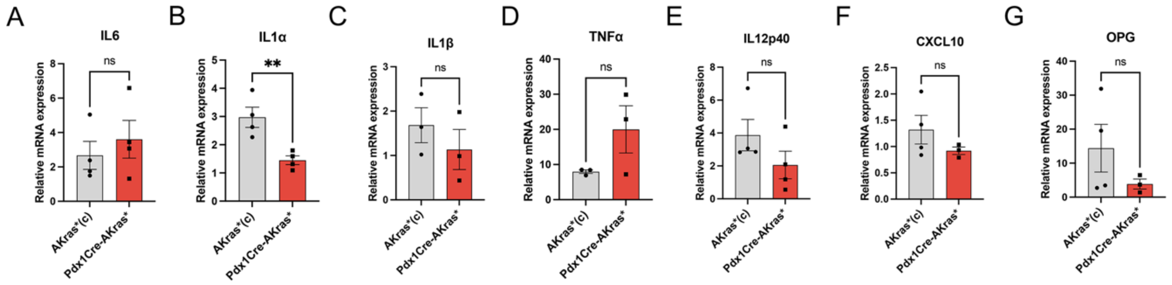


**Fig S5. Hippocampal proinflammatory gene expression is not significantly altered in Pdx1Cre-AKras* mice.**

A) RT-qPCR analysis of expression levels of IL6. Y-axis data is the relative mRNA expression ratio of AKras*(c) and Pdx1Cre-AKras* mice, and data of Pdx1Cre-AKras* mice is standardized to 1. P values were determined using two-tailed unpaired student's t-test. n=4 for each group. B) RT-qPCR analysis of expression levels of IL1α. Y-axis data is the relative mRNA expression ratio of AKras*(c) and Pdx1Cre-AKras* mice, and data of Pdx1Cre-AKras* mice is standardized to 1. P values were determined using two-tailed unpaired student's t-test. n=4 for each group. C) RT-qPCR analysis of expression levels of IL1β. Y-axis data is the relative mRNA expression ratio of AKras*(c) and Pdx1Cre-AKras* mice, and data of Pdx1Cre-AKras* mice is standardized to 1. P values were determined using two-tailed unpaired student's t-test. n=4 for each group. D) RT-qPCR analysis of expression levels of TNFα. Y-axis data is the relative mRNA expression ratio of AKras*(c) and Pdx1Cre-AKras* mice, and data of Pdx1Cre-AKras* mice is standardized to 1. P values were determined using two-tailed unpaired student's t-test. n=3 for each group. E) RT-qPCR analysis of expression levels of IL12p40. Y-axis data is the relative mRNA expression ratio of AKras*(c) and Pdx1Cre-AKras* mice, and data of Pdx1Cre-AKras* mice is standardized to 1. P values were determined using two-tailed unpaired student's t-test. n=4 for each group. F) RT-qPCR analysis of expression levels of CXCL10. Y-axis data is the relative mRNA expression ratio of AKras*(c) and Pdx1Cre-AKras* mice, and data of Pdx1Cre-AKras* mice is standardized to 1. P values were determined using two-tailed unpaired student's t-test. n=4 AKras*(c) mice and n=3 Pdx1Cre-AKras* mice. G) RT-qPCR analysis of expression levels of OPG. Y-axis data is the relative mRNA expression ratio of AKras*(c) and Pdx1Cre-AKras* mice, and data of Pdx1Cre-AKras* mice is standardized to 1. P values were determined using two-tailed unpaired student's t-test. n=4 AKras*(c) mice and n=3 Pdx1Cre-AKras* mice.

***Supplementary Table 1.*** *List of primer sets used in this study.*

| Genotyping Primers | | |
|---|---|---|
| **Target** | **Fw** | **Rv** |
| ***Tg$^{(Pdx1\text{-}cre)6Tuv/J}$*** | CTGCCACGACCAAGTGACAGC | CTTCTCTACACCTGCGGTGCT |
| **Actb$^{tm1(KrasG12D)Arge}$** | TGTATGGGATCTGATCTGGGGCCTCG | CTCGGCAGGAGCAAGGTGAGATG |
| ***Gt(ROSA)26Sor$^{tm9(CAG\text{-}tdTomato)hZE}$*** | AAGGGAGCTGCAGTGGAGTA | CCGAAAATCTGTGGGAAGTC |
| **Real-Time PCR Primers** | | |
| ***OPG*** | GTTTGCCTGGGACCAAAGTG | CACAGAGGTCAATGTCTTGGATG |
| ***TNFa*** | AGCCCACGTCGTAGCAAAC | ACAAGGTACAACCCATCGGC |
| ***IL1b*** | TGCCACCTTTTGACAGTGATG | AAGGTCCACGGGAAAGACAC |
| ***IL1a*** | CGCTTGAGTCGGCAAAGAAAT | AAGGTGCTGATCTGGGTTGG |
| ***IL12p40*** | CACCTGTGACACGCCTGAAG | CTCAGAGTCTCGCCTCCTTT |
| ***Cxcl10*** | TCCTTGTCCTCCCTAGCTCA | ATAACCCCTTGGGAAGATGG |
| ***IL6*** | GAGAGGAGACTTCACAGAGGATACC | TCATTTCCACGATTTCCCAGAGAAC |
| ***Itgam*** | ACGCAGACAGGAAGTACCAG | CCCAGCAAGGGACCATTAGAG |
| ***Cd68*** | CTAGGACCGCTTATAGCCCAAG | GTGAAGGATGGCAGGAGAGT |
| ***MertK*** | AGGAAGGTAGGTCCTGGGTG | TCTCGGCAGTGCCTAAGGAT |

***Supplementary Table 2***. *List of Antibodies used in this study.*

| Antigen | Clone | Source | Cat. Number | Dilution |
|---|---|---|---|---|
| DCX | Mouse monoclonal | Santa-Cruz | sc271390 | 1/100 |
| GFAP | Chicken polyclonal | Abcam | ab1674 | 1/500 |
| SOX2 | Rabbit polyclonal | Proteintech | 11064 | 1/200 |
| PDX1 | Guinea pig polyclonal | Abcam | ab47308 | 1/150 |
| BrdU | Rat monoclonal | Abcam | ab6326 | 1/200 |
| Anti-Mouse IgG - Alexa Fluor 488 | Goat polyclonal | Invitrogen | a11001 | 1/500 |
| Anti-Rabbit IgG - Alexa Fluor 568 | Goat polyclonal | Invitrogen | A11011 | 1/500 |
| Anti-Chicken IgY - Alexa Fluor 568 | Goat polyclonal | Abcam | ab175477 | 1/500 |
| Anti-Rat IgG - CF488A | Goat polyclonal | Biotium | 20096 | 1/500 |

***Supplementary Table 3.*** *Concentration levels of measured cytokines (ng/ml).*

| | **GM-CSF** | **IFNγ** | **IL4** | **IL17F** |
|---|---|---|---|---|
| **Akras*(c) - P40** | 0.005230453 | 0.016760402 | 0.013747599 | 0.006945588 |
| **Pdx1Cre-AKras* - P40** | 0.029329303 | 0.174900144 | 0.041869717 | 0.013059257 |
| **Akras*(c) - P21** | 0.005230453 | 0.016760402 | 0.013747599 | 0.007755183 |
| **Pdx1Cre-AKras* - P21** | 0.005230453 | 0.016760402 | 0.00494971 | 0.035887082 |
| **Xeno$^{Sham}$(c)** | 0.005230453 | 0.016760402 | 0.013747599 | 0.015653677 |

| | | | | |
|---|---|---|---|---|
| **Xeno$^{Panc-1}$ - Low Burden** | 0.005230453 | 0.016760402 | 0.013747599 | 0.00521245 |
| **Xeno$^{Panc-1}$ - High Burden** | 0.001083635 | 0.015735066 | 0.00186115 | 0.006945588 |

| | **IL1α** | **CXCL10** | **MCP-1 (CCL2)** | **TNFα** |
|---|---|---|---|---|
| **Akras*(c) - P40** | 0.00194193 | 0.040834788 | 0.102083866 | 0.006242341 |
| **Pdx1Cre-AKras* - P40** | 0.049890134 | 0.445542576 | 1.056231186 | 0.095668376 |
| **Akras*(c) - P21** | 0.00194193 | 0.107581464 | 0.470618767 | 0.006242341 |
| **Pdx1Cre-AKras* - P21** | 0.00194193 | 0.169195666 | 0.705281918 | 0.006242341 |
| **Xeno$^{Sham}$(c)** | 0.00194193 | 0.148678221 | 0.646720511 | 0.006242341 |
| **Xeno$^{Panc-1}$ - Low Burden** | 0.00098656 | 0.28371351 | 0.968920871 | 0.006242341 |
| **Xeno$^{Panc-1}$ - High Burden** | 0.000988257 | 0.284712823 | 5.663838513 | 0.006242341 |

| | **IL22** | **IL1β** | **TNFSF11** | **RANK** |
|---|---|---|---|---|
| **Akras*(c) - P40** | 0.015226337 | 0.022862369 | 0.003976518 | 0.829506238 |
| **Pdx1Cre-AKras* - P40** | 0.083269865 | 0.824530343 | 0.041440127 | 2.774256631 |
| **Akras*(c) - P21** | 0.015226337 | 0.022862369 | 0.004985409 | 1.16676474 |
| **Pdx1Cre-AKras* - P21** | 0.013133865 | 0.036672264 | 0.026015649 | 1.934970377 |
| **Xeno$^{Sham}$(c)** | 0.004533432 | 0.022862369 | 0.003882301 | 0.445215583 |
| **Xeno$^{Panc-1}$ - Low Burden** | 0.009258255 | 0.016726272 | 0.004678385 | 0.871568752 |
| **Xeno$^{Panc-1}$ - High Burden** | 0.015226337 | 0.001236021 | 0.004942156 | 1.963626935 |

| | **IL10** | **IL6** | **G-CSF** | **IL17A** |
|---|---|---|---|---|
| **Akras*(c) - P40** | 0.028303698 | 0.022862369 | 0.016238226 | 0.007773205 |
| **Pdx1Cre-AKras* - P40** | 0.129132487 | 0.097298177 | 0.069341886 | 0.039134993 |
| **Akras*(c) - P21** | 0.028302698 | 0.022862369 | 0.016238226 | 0.007773205 |
| **Pdx1Cre-AKras* - P21** | 0.028302698 | 0.022862369 | 0.002893678 | 0.00413108 |
| **Xeno$^{Sham}$(c)** | 0.028302698 | 0.022862369 | 0.009007956 | 0.007773205 |
| **Xeno$^{Panc-1}$ - Low Burden** | 0.028302698 | 0.022862369 | 0.016238226 | 0.007773205 |
| **Xeno$^{Panc-1}$ - High Burden** | 0.028302698 | 0.342208466 | 0.099248522 | 0.007773205 |

| | **OPG** | **M-CSF** | **IL12p40** |
|---|---|---|---|
| **Akras*(c) - P40** | 0.93893785 | 0.009253577 | 0.022862369 |
| **Pdx1Cre-AKras* - P40** | 8.590224749 | 0.049017904 | 0.616685528 |
| **Akras*(c) - P21** | 5.225163658 | 0.012833146 | 0.046153107 |
| **Pdx1Cre-AKras* - P21** | 4.29605645 | 0.058032697 | 0.469421435 |
| **Xeno$^{Sham}$(c)** | 2.368998877 | 0.002816998 | 0.022862369 |
| **Xeno$^{Panc-1}$ - Low Burden** | 3.990403653 | 0.006154101 | 0.022862369 |
| **Xeno$^{Panc-1}$ - High Burden** | 7.323080054 | 0.00419868 | 0.022862369 |
